# A dominant-negative avirulence effector of the barley powdery mildew fungus provides mechanistic insight to barley MLA immune receptor activation

**DOI:** 10.1101/2023.01.11.523539

**Authors:** Emma E Crean, Merle Bilstein-Schloemer, Takaki Maekawa, Paul Schulze-Lefert, Isabel ML Saur

## Abstract

Nucleotide-binding leucine-rich repeat receptors (NLRs) recognize pathogen effectors to mediate plant disease resistance, which is often accompanied by a localized host cell death response. Effectors can escape NLR recognition through various polymorphisms, allowing the pathogen to proliferate on previously resistant host plants. The powdery mildew effector AVR_A13_-1 is recognized by the barley NLR MLA13 and activates host cell death. We demonstrate here that a virulent form of AVR_A13_, called AVR_A13_-V2, escapes MLA13 recognition by substituting a serine for a leucine residue at the C-terminus. Counterintuitively, this substitution in AVR_A13_-V2 resulted in an enhanced MLA13 association and prevented the detection of AVR_A13_-1 by MLA13. Therefore, AVR_A13_-V2 is a dominant-negative form of AVR_A13_ and has likely contributed to the breakdown of *Mla13* resistance. Despite this dominant-negative activity, AVR_A13_-V2 failed to suppress host cell death mediated by the MLA13 auto-active “MHD” variant. Neither AVR_A13_-1 nor AVR_A13_-V2 interacted with the MLA13 auto-active variant, implying that the binding moiety in MLA13 that mediates association with AVR_A13_-1 is altered after receptor activation. We also show that mutations in the MLA13 coiled-coil signalling domain, which were thought to impair Ca^2+^-channel activity and NLR function, instead resulted in MLA13 auto-active cell death. The data constitute an important step to define intermediate receptor conformations during NLR activation.

## Introduction

During infection of their host, pathogens secrete numerous molecules that act either extracellularly or inside host cells. Some of these molecules act as virulence factors (so-called effectors) to manipulate the host’s physiology in favour of the pathogen. Disease resistance of a plant against a pathogen is often mediated by resistance genes encoding nucleotide-binding leucine-rich repeat receptors (NLRs) (Maekawa *et al*., 2011b; Jones *et al*., 2016). NLRs recognize effectors by direct binding or by indirectly detecting effector-mediated alterations of host targets (guardees) or their mimics (decoys) (Cesari, 2018). Effector-mediated NLR activation is often linked to localized host cell death (Dodds and Rathjen, 2010; Saur and Hückelhoven, 2021; Maekawa *et al*., 2022) and recognized effectors are called avirulence (AVR) effectors. Diversification of genes encoding AVRs can lead to loss of recognition by the respective NLR, resulting in pathogen virulence and breakdown of disease resistance (Märkle *et al*., 2022). In case of direct AVR recognition, the NLR can usually no longer bind the diversified effector proteins of virulent pathogen isolates (Saur *et al*., 2021).

NLRs are modular multidomain proteins with a central NB (nucleotide-binding) domain and C-terminal leucine-rich repeats (LRRs). At the N-terminus, most NLRs encode either a Toll/Interleukin-1 receptor-like (TIR) or a coiled-coil (CC) domain, classifying the majority of NLRs into either TIR-type NLRs (TNLs) or CC-type NLRs (CNLs) (Shao *et al*., 2016). A subgroup of CNLs (also called RPW8-like NLRs or RNLs) are the helper NLRs NRG1 (N REQUIREMENT GENE 1) and ADR1 (ACTIVATED DISEASE RESISTANCE GENE 1) that are genetically required for TNL-mediated disease resistance (Saile *et al*., 2020). The N-terminal CC and TIR domains mediate NLR signal emission upon NLR activation (Swiderski *et al*., 2009; Bernoux *et al*., 2011; Collier *et al*., 2011; Maekawa *et al*., 2011a; Williams *et al*., 2014). In the absence of matching pathogen effectors, CC and TIR domains are locked in inactive conformations and this auto-inhibition is mediated by inter-domain interactions between the N-terminal domains with the NB and LRR domains (Burdett *et al*., 2019; Saur *et al*., 2021; Tamborski *et al*., 2022). Although structural information on intermediate forms between inactive and active signalling NLRs is limited to the structure of the *Arabidopsis thaliana* CNL ZAR1 (HOPZ-ACTIVATED RESISTANCE 1) (Wang *et al*., 2019b), NLR activation is thought to be a multistep process (Förderer *et al*., 2022b). The first activation step is ligand binding, which induces a steric clash between the LRR and the NB domain. The resulting open conformation of the NB domain then allows exchange of ADP (inactive) to ATP (active), which in turn induces allosteric changes to release the conformational auto-inhibition of the CC or TIR domains. This induces NLR oligomerization and these NLR oligomers are referred to as resistosomes (Förderer *et al*., 2022b). Certain amino acid replacements within the conserved MHD motif of the NB domain mimic ATP binding and thus result in an active NLR conformation (Dinesh-Kumar and Baker, 2000; Bendahmane *et al*., 2002; Paulmurugan *et al*., 2002; Howles *et al*., 2005; Gao *et al*., 2011; Bai *et al*., 2012; Ntoukakis *et al*., 2013, 2014; Roberts *et al*., 2013; Nishimura *et al*., 2017). The N-terminal portion of the LRR domain in CNLs also contributes to receptor auto-regulation through interactions with CC and NB domains and amino acid exchanges at these sites can affect NLR auto-activity (Rairdan and Moffett, 2006; Slootweg *et al*., 2013; Burdett *et al*., 2019; Förderer *et al*., 2022a; Tamborski *et al*., 2022). For receptor activation *via* direct effector recognition, amino acids in the LRR can have additional functions as effector contact sites and can define the specificity of effector recognition (Jia *et al*., 2000; Shen *et al*., 2003; Dodds *et al*., 2006; Bauer *et al*., 2021; Tamborski *et al*., 2022; Förderer *et al*., 2022b). Upon direct effector recognition by the LRR or other integrated domains, effector binding correlates directly with NLR signal activation and studies on the *Magnaporthe oryzae* effectors AvrPik and AVR-Pia and the rice NLRs Pik and RGA5, respectively, argue for an affinity threshold between receptor and effector for activation of NLR immune signalling and pathogen resistance (Ortiz *et al*., 2017; de la Concepcion *et al*., 2018).

While the mechanisms underlying the restriction of pathogen growth by resistosomes is not fully elucidated, recent cryo-EM structures of multiple resistosomes (Wang *et al*., 2019a, *b*; Ma *et al*., 2020; Martin *et al*., 2020; Förderer *et al*., 2022a) revealed fundamental differences in immune signalling initiated by TNLs and CNLs: the pentameric resistosomes of *A. thaliana* ZAR1 CNL and wheat Sr35 CNL have calcium ion-permeable non-selective cation channel activity (Bi *et al*., 2021; Förderer *et al*., 2022a). The funnel-shaped ZAR1 cation channel is formed by the N-terminal CC domain α1-helix of the ZAR1 resistosome (Wang *et al*., 2019a, *b*). Substitutions of negatively charged amino acids to alanine in the inner lining of the funnel abolishes Ca^2+^ channel and cell death activity and ZAR1-mediated resistance (Wang *et al*., 2019b; Bi *et al*., 2021). The α1-helix region of the wheat Sr35 resistosome is not well resolved in the cryo-EM structure and Sr35 α1-helix amino acid exchanges equivalent to those in ZAR1 do not affect AvrSr35-dependent Sr35 resistosome channel and cell death activity (Förderer *et al*., 2022a; Zhao *et al*., 2022), suggesting differences in Ca^2+^ signalling functions between ZAR1 and Sr35 resistosomes. Effector binding to the TNLs RPP1 (RECOGNITION OF PERONOSPORA PARASITICA 1) and ROQ1 (RECOGNITION OF XopQ 1) from *A. thaliana* and *Nicotiana benthamiana*, respectively, induces the formation of homotetrameric complexes stimulating TIR enzyme activity. The resistosome TIR enzyme, but also TIR-only proteins, produce a variety of nucleotide-based second messenger molecules (Horsefield *et al*., 2019; Wan *et al*., 2019; Yu *et al*., 2022; Huang *et al*., 2022; Jia *et al*., 2022), some of which serve as ligands to activate the EDS1 protein family plus the signalling/helper CNLs ADR1 or NRG1 (Lapin *et al*., 2019; Huang *et al*., 2022; Jia *et al*., 2022). ADR1 and NRG1 can also function as calcium ion-permeable nonselective cation channels (Jacob *et al*., 2021), and as such disruption of Ca^2+^ homeostasis appears to be central in CNL and TNL resistosome signalling.

The polymorphic barley *Mildew locus A (Mla)* encodes allelic variants of CNLs (MLA NLRs), each conferring isolate-specific disease resistance to the barley powdery mildew fungus *Blumeria graminis* f. sp. *hordei (Bgh)* (Moseman and Schaller, 1960; Glawe, 2008; Seeholzer *et al*., 2010; Maekawa *et al*., 2019). Some barley MLA receptors and *Mla* homologs confer additional resistance to isolates of unrelated fungal pathogens (Periyannan *et al*., 2013; Mago *et al*., 2015; Chen *et al*., 2017; Bettgenhaeuser *et al*., 2021; Ortiz *et al*., 2022; Brabham *et al*., 2022). The *Bgh* effectors recognized by barley MLAs are known as AVR_A_ effectors (Moseman and Schaller, 1960; Jorgensen, 1994) and diversified variants that have escaped *Mla* recognition are designated as AVR_A_-V variants (Lu *et al*., 2016). To date, full length structures of inactive or effector-activated MLAs are not available, but protein interaction assays suggest a direct interaction between at least some MLA NLRs and matching AVR_A_ effectors (Saur *et al*., 2019a). Most amino acids under positive selection of *Mla* resistance specificities map to the predicted solvent-exposed sites of the LRR, suggesting that these serve as AVR_A_ contact residues (Seeholzer *et al*., 2010; Maekawa *et al*., 2019), but interaction between effectors and MLA LRR domain deletion constructs could not be shown. Most of the known *Bgh* AVR_A_ effectors are sequence-unrelated, but share a common fold reminiscent of ribonucleases lacking catalytic residues (Bauer *et al*., 2021).

*Mla13* in barley confers resistance to most *Bgh* isolates representing a global pathogen population because these avirulent isolates express recognised AVR_A13_-1/BLGH_02099 (Lu *et al*., 2016; Saur *et al*., 2019a). AVR_A13_-1 is directly recognized by MLA13 and effector recognition drives MLA13-mediated cell death upon transient co-expression of *Mla13* and *AVR_a13_-1* in barley protoplasts and heterologous *N. benthamiana* leaves. *AVR_a13_-1/BLGH_02099* is polymorphic in the *Mla13*-virulent *Bgh* isolates CC52 and B103 and the resulting gene products are named AVR_A13_-V1 and AVR_A13_-V2, respectively (Lu *et al*., 2016). AVR_A13_-V1 represents a truncated version of AVR_A13_-1, and after transient gene overexpression *in planta*, the AVR_A13_-V1 protein is unstable and often not detectable. Currently, it cannot be ruled out that the lack of AVR_A13_-V1 recognition by MLA13 is solely due to AVR_A13_-V1 protein instability (Saur *et al*., 2019a). AVR_A13_-V2 carries five C-terminal amino acids (VRATL) that correspond to eight unrelated amino acids in AVR_A13_-1 (TCMVSSPE). Not in agreement with the virulent pathotype of *Bgh* isolate B103 on *Mla13* barley, interaction assays *in planta* and in yeast indicated a stable association between AVR_A13_-V2 and MLA13 (Saur *et al*., 2019a).

Because receptor-effector interaction is commonly linked to receptor activation, we aimed here to investigate the seeming paradox of MLA13 inactivity despite stable AVR_A13_-V2 - MLA13 association By applying proximity-dependent protein labelling (BioID), yeast-2-hybrid (Y2H) interaction assays and structural prediction (Alphafold2) in combination with *in planta* expression of naturally occurring AVR_A13_ effector variants and by generating deletion and hybrid constructs, we demonstrate that a single surface-exposed amino acid at the C-terminus of AVR_A13_ effectors determines the association with and activation of MLA13. Our data also reveal that AVR_A13_-V2 acts as dominant-negative effector on MLA13-mediated cell death. This proposes that breakdown of *Mla13*-mediated resistance can be explained by *Bgh* isolates carrying dominant-negative AVR_A13_-V2. We also demonstrate that amino acid exchanges in the MLA13 NB and LRR domains compromise effector binding. In turn, amino acid changes in the MLA13 CC domain predicted to disrupt cation channel activity, do not affect MLA13-mediated cell death. Nevertheless, inhibition of Ca^2+^ and other cation channels by LaCl3 impaired MLA13-mediated cell death of barley protoplasts. Collectively, these results provide insights and tools for understanding the conformational changes NLRs undergo during effector-mediated NLR resistosome activation.

## Results

### The C-terminus of AVR_A13_ effectors determines interaction with and activation of MLA13

The C-terminally located polymorphisms between genes encoding avirulent AVR_A13_-1 effector (activates MLA13-specified cell death) and virulent AVR_A13_-V1 or AVR_A13_-V2 variants (unable to activate MLA13 cell death; Fig. 1A) indicate a role of the AVR_A13_-1 C-terminus in the interaction with and activation of MLA13. Previously, no avirulence activity could be detected for AVR_A13_-V1, but this could be attributed to its protein instability upon transient expression *in planta* (Lu *et al*., 2016; Saur *et al*., 2019a). Here we aimed to stabilize AVR_A13_-V1 protein by fusion with an epitope to retest the association patterns of the AVR_A13_ variants with MLA13 *in planta*. To this end, we fused the three effector variants to a biotin ligase (BirA), and indeed this fusion allowed immunodetection of the AVR_A13_-V1 protein at levels comparable to the other two variants in *N. benthamiana* leaves (Fig. S1). We also confirmed the functionality of the tagged proteins by demonstrating MLA13-specified cell death induced by AVR_A13_-1-BirA-4xMyc (Fig. S1). We detected biotinylated MLA13, but not MLA1 or MLA7 protein, in samples expressing *Mla13-4Myc* together with *AVR_a13_-1-BirA* or *AVR_a13_-V2-BirA*, but not *AVR_a13_-V1-BirA* after biotin treatment followed by a streptavidin pull-down (Fig. S1B). Given that AVR_A13_-V1 lacks the 42 C-terminal amino acids of AVR_A13_-1 (Fig. 1A), the data provides experimental evidence that the C-terminal half of AVR_A13_ is needed for the association and activation of MLA13 receptor.

**Fig. 1:**
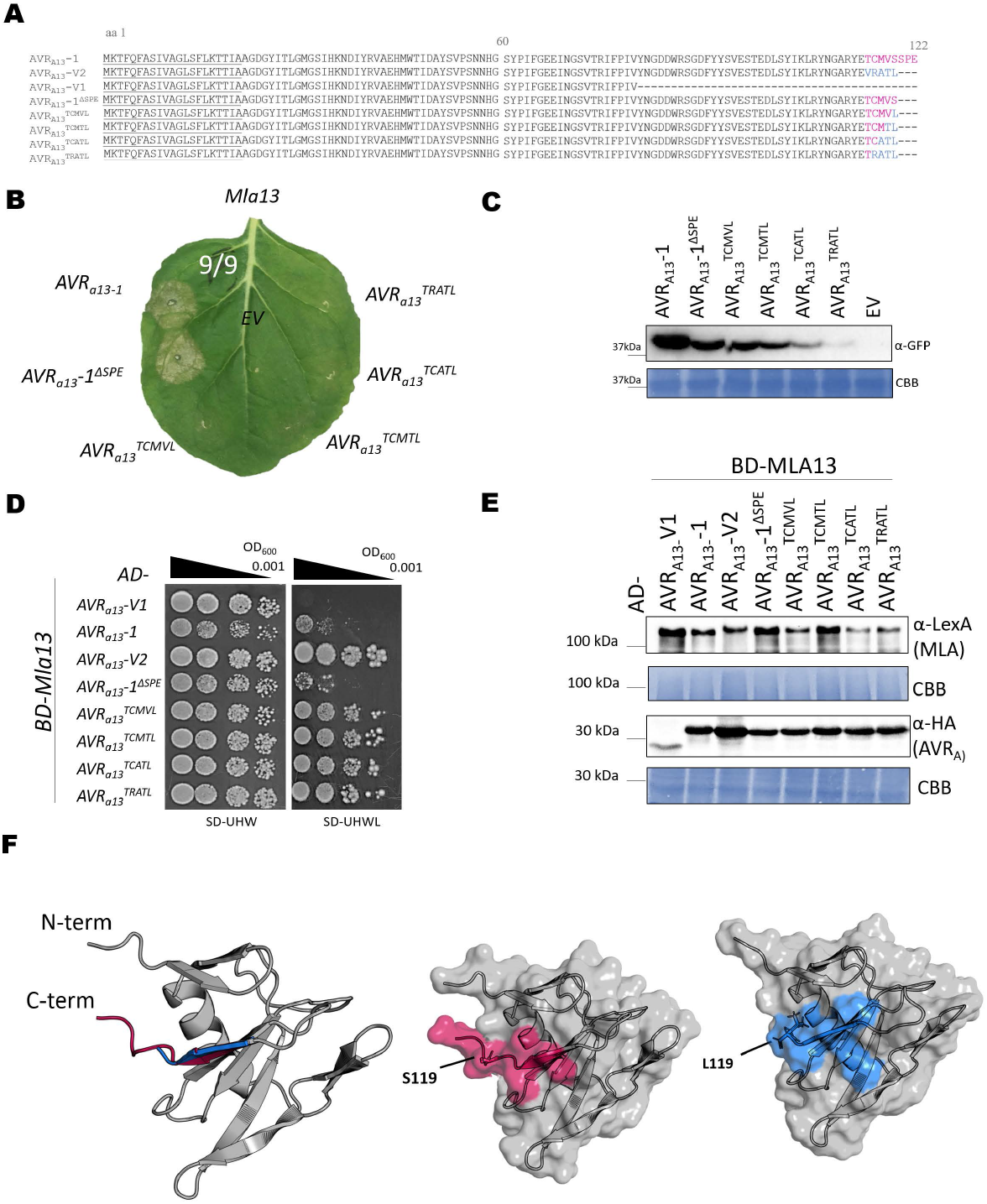
The C-terminus of AVR_A13_ effectors controls interaction with and activation of MLA13. **(A)** Amino acid (aa) alignment of AVR_A13_ variants analysed for interaction with MLA13 and inhibition of Mla13-mediated cell death. Signal peptide (SP) residues are underlined; aa in pink and blue highlight the aa variation between AVR_A13_-V2 and AVR_A13_-1, respectively. **(B, C)** *Nicotiana benthamiana* leaves were transformed transiently with *35S:Mla13-4Myc* (pGWB517) with one of the *AVR_a13_* variants lacking SPs cloned between the 35S promoter and a C-terminal *mYFP* sequence or *empty vector* (*EV*). **(B)** Cell death was determined three days post transformation and figures shown are representatives of at least nine independent leaves from at least three independent plants. **(C)** Protein stability of the AVR_A13_ variants fused to mYFP corresponding to constructs of B. Leaf tissue was harvested two days post infiltration. Total protein was extracted, separated by gel electrophoresis and probed by anti-GFP western blotting (WB). **(D, E)** Yeast cells were co-transformed with *Mla13* fused N-terminally to the *LexA* binding domain sequence (BD) and *AVR_a13_* variants lacking SPs fused N-terminally to the *B42* activation domain (AD) and 1xHA tag sequence as indicated. Growth of transformants was determined on selective growth media containing raffinose and galactose as carbon sources but lacking uracil, histidine and tryptophan (-UHW), and interaction of proteins was determined by leucine reporter activity reflected by growth of yeast on selective media containing raffinose and galactose as carbon sources but lacking uracil, histidine, tryptophan and leucine (-UHWL). Figures shown are representatives of at least three experiments and pictures were taken 6 to 8 days after drop out. (**E**) Protein levels of BD-MLA13 and AD-AVR_A_ variants corresponding to yeast of D. Yeast transformants were grown in raffinose and galactose containing selective media lacking uracil, tryptophan, and histidine to OD_600_ = 1. Then, cells were harvested, total protein extracted, separated by gel electrophoresis, and western blots (WB) were probed with anti-LexA or anti-HA antibodies as indicated. CBB: Coomassie brilliant blue. (**F**) Cartoon and surface representations for the top rank model of AVR_A13_-1 and AVR_A13_-V2 from AlphaFold2 (pLDDT_overall_= 89, pLDDT_L/S119_ >80). Residues highlighted in pink correspond to the AVR_A13_-1 C-terminal residues and those in blue correspond to the AVR_A13_-V2 C-terminal residues.

Both, AVR_A13_-1 and AVR_A13_-V2 associate with MLA13, but only AVR_A13_-1 activates MLA13-mediated cell death (Saur *et al*., 2019a) (Fig. S1). To delineate the AVR_A13_-1 amino acids required for MLA13 cell death activation, we generated a truncated AVR_A13_-1 construct (AVR_A13_-1^ΔSPE^) and four hybrid variants of AVR_A13_-1 and AVR_A13_-V2, which differ from AVR_A13_-1^ΔSPE^ by one, two, three and four C-terminal amino acids, respectively (Fig. 1A). We then measured the ability of AVR_A13_-1^ΔSPE^ and the hybrid variants to induce MLA13-mediated cell death upon transient expression in *N. benthamiana* leaves (Fig. 1B and 1C). AVR_A13_-1^ΔSPE^ was the only engineered construct that induced MLA13-specified cell death comparable to AVR_A13_-1 in these assays (Fig. 1B). Therefore, the three C-terminal amino acids are dispensable for the avirulence activity of AVR_A13_-1. The data demonstrates that the replacement of serine to leucine at position 119 abrogated MLA13-mediated cell death in *N. benthamiana*, suggesting that this serine in AVR_A13_-1 and AVR_A13_-1^ΔSPE^ is crucial for cell death activation (Fig. 1B).

MLA13 interacts more efficiently with AVR_A13_-V2 than with AVR_A13_-1, and this enhanced association correlates with the inability to induce MLA13-mediated cell death (Saur *et al*., 2019a). We therefore tested the association of AVR_A13_-1^ΔSPE^ and the AVR_A13_-1/AVR_A13_-V2 hybrid variants with MLA13. Protein stability of AVR_A13_ hybrid variants varies *in planta*, which makes the assessment of quantitative differences of the interactions difficult (Fig. 1C). We then used a Y2H assay drop out series to evaluate putative quantitative differences, as in this system the corresponding prey and bait variants accumulated to comparable levels. We fused *Mla* N-terminally to the *LexA* binding domain sequence (*BD-Mla13*) and the *AVR_a13_* variant genes to the *B42* activation domain (*AD-AVR_a13_*) and determined yeast growth in the absence of leucine as a proxy for protein interaction. We found that yeasts co-expressing *BD-Mla13* with the *AD-AVR_a13_-1* and *AD-AVR_a13_-1^ΔSPE^* grew less in the dilution series than yeasts carrying *AD-AVR_a13_-V2* or any of the *AD-AVR_a13_* hybrid constructs (Fig. 1D). No growth was detected when *BD-Mla13* was co-expressed with *AD-AVRa13-V1* or when any *AVR_a13_* variant was co-expressed with *BD-Mla1* (Fig. 1D and Fig. S2). The data imply that L^119^ of AVR_A13_-V2 (Fig. 1A) is responsible for the enhanced interaction with MLA13. The corresponding residue in AVR_A13_-1 is a serine. We generated structural predictions of the AVR_A13_ variants (lacking the respective signal peptides (SP)) using AlphaFold2 (pLDDT_overall_ =89, pLDDT_L/S119_ = >80) to determine if either of these residues are surface exposed. Indeed both, L^119^ of AVR_A13_-V2 ΔSP and S^119^ of AVR_A13_-1 ΔSP appear to be surface exposed in these structural models, suggesting that they are accessible for binding to MLA13 (Fig. 1F).

### AVR_A13_-V2 can act as dominant-negative effector on MLA13-mediated cell death

The observed enhanced association between MLA13 and AVR_A13_-V2 could affect *Mla13* disease resistance and possibly the activity of other MLA NLRs with resistance specificity to *Bgh*. To test this, we measured AVR_A_-induced MLA-mediated cell death in the presence of AVR_A13_-V2. Co-expression of *Mla13-4xMyc* together with *AVR_a13_-1-mYFP* (monomeric YFP) and an *empty vector (EV)* in *N. benthamiana* leaves resulted in a cell death response within 50 to 72 hours post infiltration and this response was not detectable when EV was exchanged for *AVR_a13_-V2-mYFP* (Fig. 2A). We also tested whether *AVR_a13_-V2-4xMyc* affects cell death mediated by *Mla1-3xHA* and *AVR_a1_-mYFP* or *Mla7-3xHA* and *AVR_a7_-2-mYFP* after transient expression of the constructs in *N. benthamiana*. We assessed the severity of cell death on a scale from 0 to 3 and found that AVR_A13_-1 and MLA13-mediated cell death was abrogated by the co-expression of AVR_A13_-V2. By contrast, cell death in samples expressing *AVR_a13_-V2* alongside *Mla1* and *AVR_a1_* or *Ml_a7_* and *AVRa7-2* were comparable to those in which *AVR_a13_-V2* was replaced by *EV* or *AVR_a13_-V1* (Fig. 2B). The specific inhibitory effect of *AVR_a13_-V2* on the MLA13 receptor (Fig. 2B) is not due to interference with MLA13 or AVR_A13_-1 protein stability, as both proteins were detectable at similar levels in samples with EV or unstable AVR_A13_-V1 as in samples co-expressing *AVR_a13_-V2* (Fig. 2C). Our data suggest that AVR_A13_-V2 has a dominant-negative effect on cell death activity specifically mediated by MLA13.

**Fig. 2:**
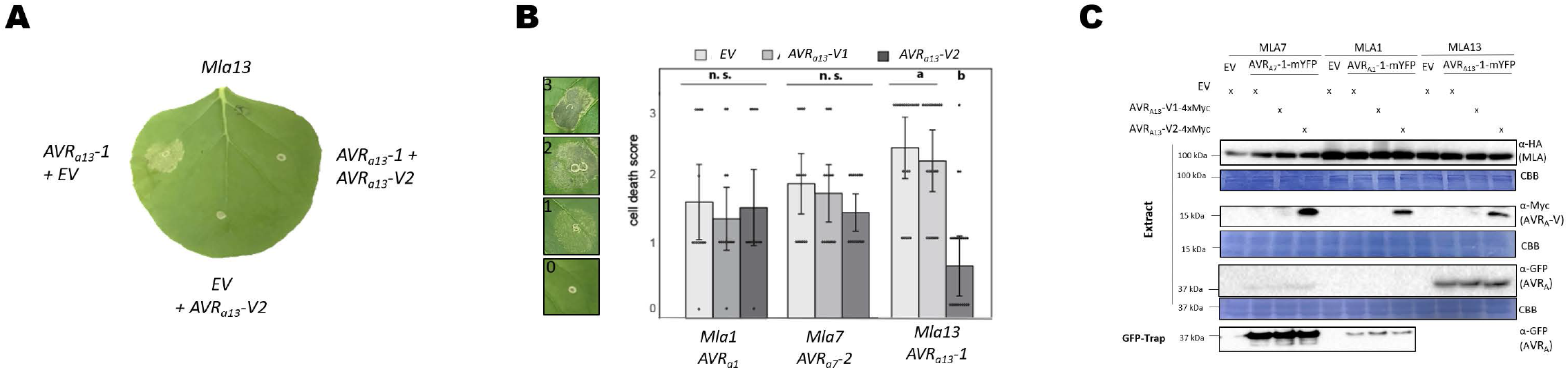
AVR_A13_-V2 can act as dominant-negative effector on MLA13. *Nicotiana benthamiana* leaves were co-transformed transiently with cDNAs of *Mla1* or *Mla7* or *MLA13* (pGWB vectors) with *AVR_a1_* or *AVR_a7_-2* or *AVR_A13_* or *empty vector (EV)* as indicated and either *AVR_A13_-V1* or *AVR-V2* or *EV* fused to epitope tags as indicated. All constructs were expressed from the 35S promoter. **(A, B)** Cell death was determined three to four days post transformation and **(B)** scored from 0 to 3 based on the cell death scale indicated. All values obtained in at least three independent experiments are indicated by dots, error bars = standard error. Differences between samples were assessed by non-parametric Kruskal-Wallis and subsequent Dunn’s tests for each MLA variant. Calculated *P* values were as follows: *Mla1: p*=0.824, *Mla7: p*=0.551 and *Mla13: p*=1.00E-06. Samples marked by identical letters in the plots do not differ significantly (p<0.05) in the Tukey test for the corresponding MLA. **(C)** Protein levels corresponding to samples of B. Leaf tissue was harvested two days post infiltration. Total protein was extracted and recovered by GFP-Trap (AVR_a1_ and AVR_a7_-2) separated by gel electrophoresis and probed by anti-HA (MLAs), anti-Myc (AVR_A13_-V2-4xMyc) or anti-GFP (AVR_a1_-mYFP, AVR_a7_-2-mYFP and AVR_A13_-1-mYFP) western blotting (WB) as indicated. CBB: Coomassie brilliant blue.

### Amino acid exchanges in the nucleotide-binding site of MLA13 compromise AVR_A13_ effector binding

Previous reports on flax TNL L6 suggest an equilibrium between inactive and active NLR conformations in the absence of pathogen effectors, but that binding of the matching effector stabilizes the active NLR conformation (Bernoux *et al*., 2016). We therefore hypothesized that avirulent AVR_A13_-1 stabilizes the active ATP-bound oligomeric conformation of MLA13. Given that AVR_A13_-V2 can inhibit MLA13-mediated cell death in the co-expression assays (Fig. 2), we hypothesized that AVR_A13_-V2 binds and stabilizes the inactive MLA13 receptor. To test this hypothesis, we applied the aforementioned Y2H approach to examine the interaction between naturally occurring AVR_A13_ variants and MLA13 variants carrying mutations in the NB domain that render the receptor inactive (P-loop mutants that cannot bind ADP or ATP at the NB domain) or auto-active (MHD mutant mimicking ATP binding at the NB domain). We first confirmed that the MLA13 P-loop mutant (MLA13^K207R^) is unable to induce cell death upon co-expression with AVR_A13_-1 (Fig. S3A) and that MLA13^D502V^ (MHD mutant) induces cell death in the absence of the matching effector (Fig. S3B), before testing their ability to bind AVR_A13_ variants in yeast. In the Y2H assay, yeast expressing BD-MLA13 together with AD-AVR_A13_-1 or AD-AVR_A13_-V2, but not AD-AVR_A13_-V1 fusion protein, grew as expected. Interestingly, none of the yeast samples co-expressing BD-MLA13^D502V^ or BD-MLA13^K207R^ together with any AVR_A13_ variants grew in the absence of leucine although all proteins were stably detectable by western blot (Fig. 3A and 3B). We observed similar results for the *Mla* homolog *Sr50*, although we detected growth of yeast expressing AD-AvrSr50 with the MHD variant Sr50^D498V^ fused N-terminally to the B42 BD (Fig. S3). However, the latter interaction was consistently weaker when compared to samples co-transformed with BD-Sr50 wild-type (WT) and AD-AvrSr50. When AD-AvrSr50 was replaced by AD-AvrSr50_QCMJC_, a variant lacking avirulence activity, no interaction was detected, which was not due to differences in protein levels between the tested effector variants (Fig. S3C and S3D).

**Fig. 3:**
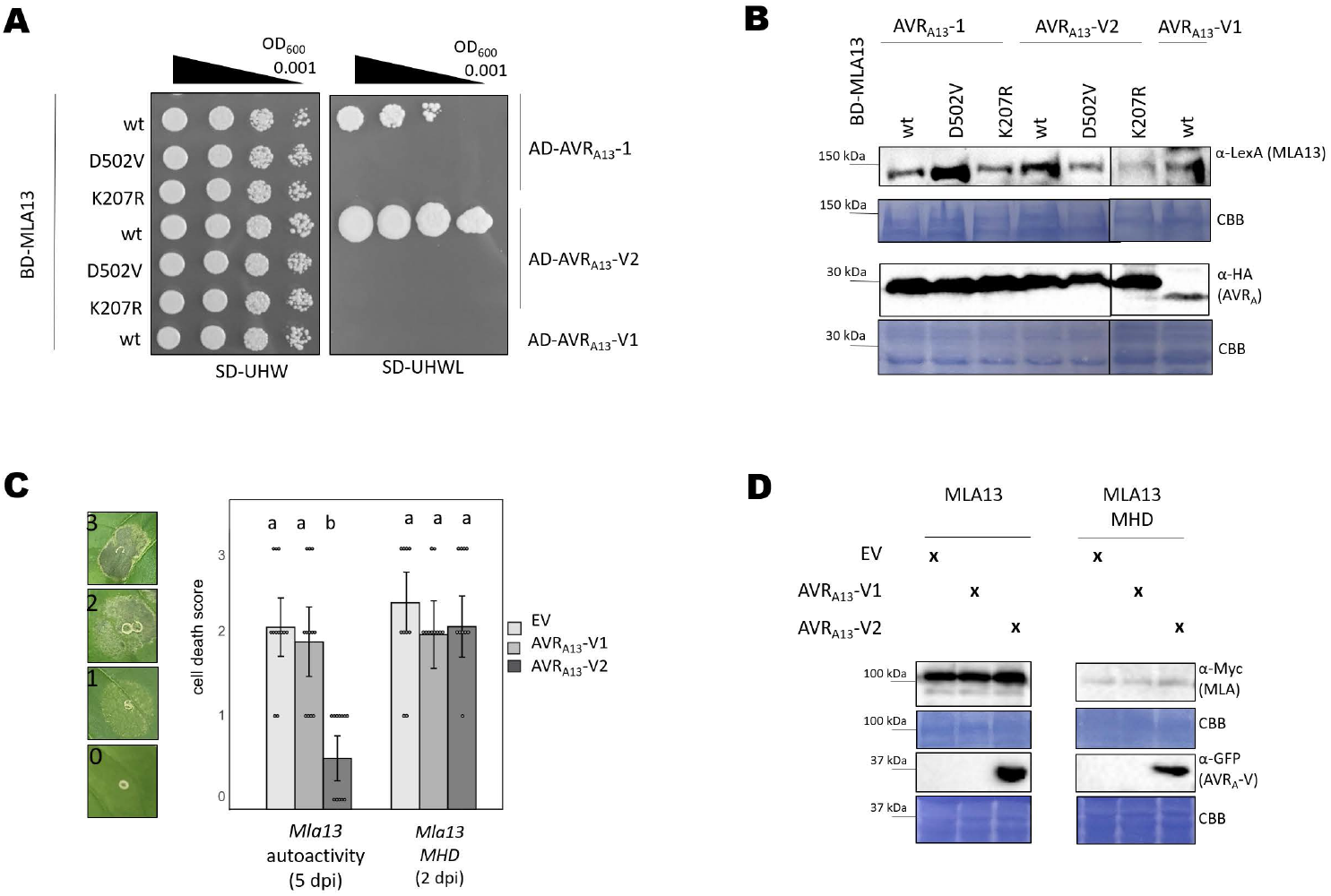
Amino acid exchanges in the nucleotide-binding site of MLA13 compromise AVRA_13_ effector binding. **(A, B)** Yeast cells were co-transformed with *Mla13 wt* or mutant variants *Mla13^D502V^* (MHD) or *Mla13 K207R* (P-loop) fused N-terminally to the *LexA* binding domain sequence (BD) and *AVR_a13_* variants lacking SPs fused N-terminally to the *B42* activation domain (AD) and *1xHA* tag sequence as indicated. **(A)** Growth of transformants was determined on selective growth media containing raffinose and galactose as carbon sources but lacking uracil, histidine and tryptophan (- UHW), and interaction of proteins was determined by leucine reporter activity reflected by growth of yeast on selective media containing raffinose and galactose as carbon sources, but lacking uracil, histidine, tryptophan and leucine (-UHWL). Figures shown are representatives of at least three experiments and pictures were taken 6 to 8 days after drop out. **(B)** Protein levels of BD-MLA13 variants and AD-AVR_A_ variants corresponding to yeast of A. Yeast transformants were grown in raffinose and galactose containing selective media lacking uracil, tryptophan, and histidine to OD_600_ = 1. Cells were harvested, total protein extracted, separated by gel electrophoresis, and western blots (WB) were probed with anti-LexA or anti-HA antibodies as indicated. **(C, D)** *Nicotiana benthamiana* leaves were co-transformed transiently with cDNAs of *AVR_a13_-V1* or *AVR_a13_-V2* or *empty vector (EV)* together with constructs encoding either MLA13 or MLA13^D502V^ (pAM-PAT vector) as indicated and under the control of the 35S promoter sequence at a 2:1 ratio. **(C)** Cell death was determined two (MLA13 MHD) to five days (MLA13) post transformation and scored from 0 to 3 based on the cell death scale indicated. All values obtained in at least three independent experiments are indicated by dots, error bars = standard deviation. Differences between samples were assessed by non-parametric Kruskal-Wallis and subsequent Dunn’s tests for each MLA variant. Calculated *P* values were as follows: MLA13: *p*=5E-05, MLA13 MHD*; p*=0.078. Samples marked by identical letters in the plots did not differ significantly (p<0.05) in the Tukey test for the corresponding MLA. **(D)** Protein levels corresponding to samples of C. Leaf tissue was harvested 36 hours post infiltration. Total protein was extracted, separated by gel electrophoresis and probed by anti-Myc (MLAs) or anti-GFP (AVR_A13_-V2)western blotting (WB) as indicated. CBB: Coomassie brilliant blue.

AVR_A13_-V2 binds specifically and strongly to wild-type MLA13 and can inhibit MLA13-specified cell death signalling, suggesting a direct link between effector binding and cell death inhibition for this association. However, AVR_A13_-V2 cannot bind auto-active MLA13^D502V^ in the Y2H assay (Fig. 3A) and we therefore speculate that it cannot inhibit MLA13^D502V^-mediated cell death. Indeed, co-overexpression of *AVR_a13_-V2* or *AVR_A13_-V1* had no effect on the average cell death score of MLA13^D502V^-induced cell death observed as early as two days post Agroinfiltration (dpi) of the respective constructs in *N. benthamiana* leaves (Fig. 3C). Four to five days after infiltration of *N. benthamiana* leaves with Agrobacteria carrying *35S:Mla13* at OD_600_=1, we also detected effector-independent cell death mediated by wild-type MLA13 (MLA13 auto-activity). This average cell death score of 2 was significantly impaired in samples co-overexpressing *AVR_a13_-V2* (average cell death score = 0.5) but not *AVR_a13_-V1* (average cell death score = 1.9). Compared with *EV* or unstable *AVR_a13_-V1*, co-expression of *AVR_a13_-V2* had no effect on the protein levels of any of the MLA13 variants used (Fig. 3D). Of note, cell death mediated by overexpression of the MLA13 CC domain (MLA13^CC^, amino acid (aa) 1-160) was not affected by AVR_A13_-V2 (Fig. S3E and S3F).

### Different affinities between MLA13 mutant variants and AVR_A13_ effectors

The lack of AVR_A13_ interaction with both inactive and active CNL MLA13 mutant variants was unexpected, as it contrasts with previous reports on flax TNL L6 and its matching effector AvrL567 (Bernoux *et al*., 2016). We therefore investigated whether this lack of effector-receptor association could be generalized to other putatively inactive or auto-active MLA13 variants (Fig. 4A). In addition to the MLA13^D502V^ and MLA13^K207A^ variants (Fig. 3), we chose the MHD mutant variant H501G, whose auto-activity in MLA10 appears to be less pronounced than that of D502V (Bai *et al*., 2012). Receptor auto-activity was also previously reported for the F99E (mutation in the CC domain) variant of MLA10 (Bai *et al*., 2012). We also included the D284A mutant (mutation in the walker A motif of the NB site, Fig. 4A) because the corresponding variant in the *A. thaliana* CNL RPM1 (RESISTANCE TO P. SYRINGAE PV MACULICOLA 1) leads to RPM1 auto-activity (Gao *et al*., 2011). By substituting negatively charged residues in the first α-helix of MLA13 to alanine (MLA13^D2A_E17A^), we aimed to generate an MLA13 resistosome that is structurally intact but impaired in immune signalling via Ca^2+^ influx (Wang *et al*., 2019a, *b*; Bi *et al*., 2021). This hypothesis is based on the observation that the replacement of negatively charged amino acids in the α1-helix of ZAR1 abrogates Ca^2+^ influx and impairs cell death activity and ZAR1 disease resistance, but not formation and membrane association of the ZAR1 resistosome (Wang *et al*., 2019a, *b*; Bi *et al*., 2021). The S902F_F935I substitutions affect residues in the 14^th^ and 15^th^ LRRs of MLA13 (Fig. 4A) and the corresponding receptor is not expected to detect AVR_A13_-1 as it is encoded by the barley line SxGP DH-47 (cross of cultivars SusPtrit and Golden Promise), which is fully susceptible to *Bgh* isolates carrying avirulent *AVR_a13_* (Bettgenhaeuser *et al*., 2021).

**Fig. 4:**
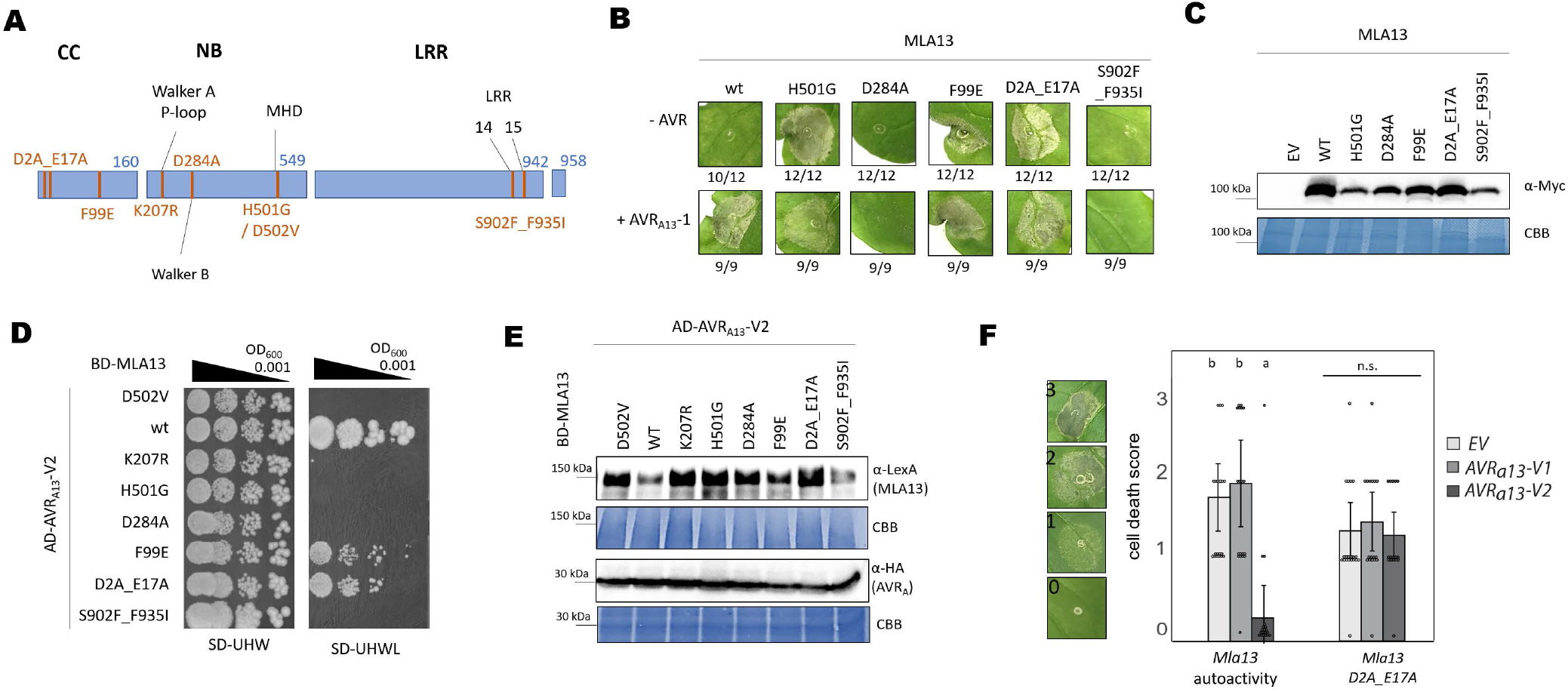
Amino acid (aa) exchanges in the coiled-coil (CC) domain de-regulate MLA13 auto-inhibition. **(A)** Amino acid (aa) changes in MLA13 mutant variants. The D2A_E17A and the F99E variants encode changes in the MLA13 coiled-coil (CC) domain, which spans from aa1 to 160. The K207R, D284A, D502V and H501G variants encode changes in the nucleotide-binding site (NB, aa 161 to 549). The S902F_F935I variants affects the leucine-rich repeats (LRR, aa 550 to 942) which are followed by a short C-terminal amino acids sequence. **(B, C)** *Nicotiana benthamiana* leaves were transformed transiently with cDNAs of one of the *Mla13* variants as indicated (pGWB517 vector) either with or without *AVR_a13_-1* lacking SPs and fused c-terminally to a *mYFP* sequence. All constructs are under the control of the 35S promotor. **(B)** Cell death was determined three days post transformation; n≥9. **(C)** Protein stability of the MLA variants fused to 4xMyc corresponding to constructs of B. Leaf tissue was harvested two days post infiltration. Total protein was extracted, separated by gel electrophoresis and probed by anti-Myc western blotting (WB) as indicated. **(D, E)** Yeast cells were co-transformed with *Mla13* variants fused N-terminally to the *LexA* binding domain (BD) sequence and *AVR_a13_-V2* lacking SPs fused N-terminally to the *B42* activation domain (AD) and *1xHA* tag sequence as indicated. Growth of transformants was determined on selective growth media containing raffinose and galactose as carbon sources but lacking uracil, histidine and tryptophan (-UHW), and interaction of proteins was determined by leucine reporter activity reflected by growth of yeast on selective media containing raffinose and galactose as carbon sources but lacking uracil, histidine, tryptophan and leucine (-UHWL). Figures shown are representatives of at least three experiments and pictures were taken 6 to 8 days after drop out. (**E**) Protein levels of BD-MLA13 variants and AD-AVR_A13_-V2 corresponding to yeast of D. Yeast transformants were grown in raffinose and galactose containing selective media lacking uracil, tryptophan, and histidine to OD_600_= 1. Then, cells were harvested, total protein extracted, separated by gel electrophoresis, and western blots (WB) were probed with anti-LexA or anti-HA antibodies as indicated. CBB: Coomassie brilliant blue. **(F)** *N. benthamiana* leaves were co-transformed transiently with cDNAs of *AVR_a13_-V1, AVR_a13_-V2* or *empty vector (EV)* together with constructs encoding the MLA13 variant as indicated and under the control of the 35S promoter sequence at a 2:1 ratio. Cell death was determined three days post transformation and scored from 0 to 3 based on the cell death scale indicated. All values obtained in at least two independent experiments are indicated by dots, error bars = standard deviation. Differences between samples were assessed by non-parametric Kruskal-Wallis and subsequent Dunn’s tests for each MLA variant. Calculated *P* values were as follows: MLA13*; p*= 9.38E-07, MLA13^D2A_E17A^: *p* = 0.77. n.s. = no significant difference.

We first tested our assumption that the MLA13 mutants exhibit altered cell death activities (inactive/auto-active) compared to wild-type. We expressed the corresponding gene constructs in *N. benthamiana* leaves and qualitatively determined cell death in the presence and absence of AVR_A13_-1. As reported for other MLA variants (Bai *et al*., 2012), MLA13^H501G^ and MLA13^F99E^ showed effector-independent cell death activity in this assay. In contrast, the Walker A-motif mutant MLA13^D284A^ and SusPtritis MLA13^S902F_F935I^ receptor variants are unable to trigger host cell death when expressed together with *AVR_a13_-1*. However, expression of MLA13^D2A_E17A^, which is thought to be impaired in Ca^2+^ and cell death signalling (Bi *et al*., 2021), resulted in effector-independent cell death in *N. benthamiana* leaves within 2 dpi (Fig. 4B). All MLA13 variants are detectable as fusion proteins after transient expression in *N. benthamiana* (Fig. 4C).

We next determined the ability of AVR_A13_-V2 to bind MLA13^H501G^, MLA13^F99E^, MLA13^D284A^, MLA13^D2A_E17A^ and MLA13^S902F_F935I^ in a Y2H assay. Again, MLA13^D502V^ and MLA13^K207R^ variants served as negative controls. Yeast samples expressing *AD_AVR_a13_-V2* together with wild-type *BD-Mla13* grew to a dilution of OD600 = 0.001 and to a dilution of OD600 = 0.01 when wild-type MLA13 was replaced with MLA13^D2A_E17A^ or MLA13^F99E^. When wild-type MLA13 was replaced by MLA13^D284A^, MLA13^K207R^ o MLA13^S902F_F935I^ showed growth in the absence of leucine (Fig. 4D) although these MLA13 variants are stably expressed in yeast (Fig. 4E). The MLA F99 residue is not conserved in other CNLs and therefore, the currently available CNL resistosome structures of ZAR1 and Sr35 cannot give functional insight to the role of this residue. However, the ZAR1 resistosome structures postulate that upon ligand binding, the release of the α1-helix in CNLs is an important conformational change that occurs immediately before resistosome formation (Wang *et al*., 2019a, *b*). We thus speculate that the auto-activity of MLA13^D2A_E17A^ is a result of mutation-induced α1-helix release. If this is the case, then this auto-activity cannot be inhibited by the dominant-negative AVR_A13_-V2 ligand. When compared to EV, co-expression of *AVR_a13_-V2-mYFP* with MLA13^D2A_E17A^ in *N. benthamiana* leaves had indeed no impact on the average cell death score, whereas auto-activity of wild-type MLA13 was again inhibited by co-expression of *AVR_a13_-V2-mYFP* (Fig. 4F).

### Activity of cation channels is required for MLA13 cell death

In ZAR1, the negatively charged residues on the inner lining of the ZAR1 resistosome funnel are required for Ca^2+^ channel activity, and substitutions of these amino acids impaired ZAR1 signalling (Bai *et al*., 2012; Wang *et al*., 2019b). By contrast, such substitutions in Sr35 had no effect on cell death or channel activity (Förderer et al., 2022a), and the same appears to be true for MLA13 D2A_E17A (Fig. 4B). The data suggest that MLA13 does not require the negatively charged amino acids of the α1-helix in the CC domain for cell death signalling. We thus aimed to determine whether Ca^2+^ channel activity is needed for MLA13-mediated cell death in barley by applying the potent cation channel inhibitor LaCl_3_. Toward this end, we expressed a luciferase (LUC) reporter together with *AVR_a13_-1* in barley mesophyll protoplasts, prepared from the *Mla13*-containing near-isogenic backcross line Manchuria (CI 16155), and measured LUC activity as an indicator of protoplast viability. Protoplasts from the cultivar Manchuria (CI 2330), which lack *Mla13*, served as control. With increasing LaCl_3_ concentration, we observed a reduction in LUC activity by up to 50% of CI 2330 protoplasts (20 μM LaCl_3_), suggesting a detrimental impact of LaCl_3_ treatment on protoplast viability independent of *Mla13* or a reduction in LUC activity independent of cell death. Nonetheless, in the absence of LaCl_3_, LUC activity is on average more than 70% lower in *Mla13* protoplasts transfected with the *AVR_a13_-1* construct than in protoplasts that do not express *Mla13* (Fig. 5A). This difference in LUC activity between the two samples diminishes with increasing LaCl_3_ concentration and is no longer significant in samples treated with 10 μM LaCl_3_. Although LUC activity decreases with increasing LaCl_3_ concentrations, LaCL_3_ treatment does not affect AVR_A13_-1 protein stability in protoplasts of the cultivar Manchuria (Fig. 5B). Although we cannot exclude that LaCl_3_ treatment affects *Mla13* expression in barley line CI 16155, our data show that blocking the function of cation channels by LaCl_3_ compromises MLA13-mediated cell death in barley leaf protoplasts.

**Fig. 5:**
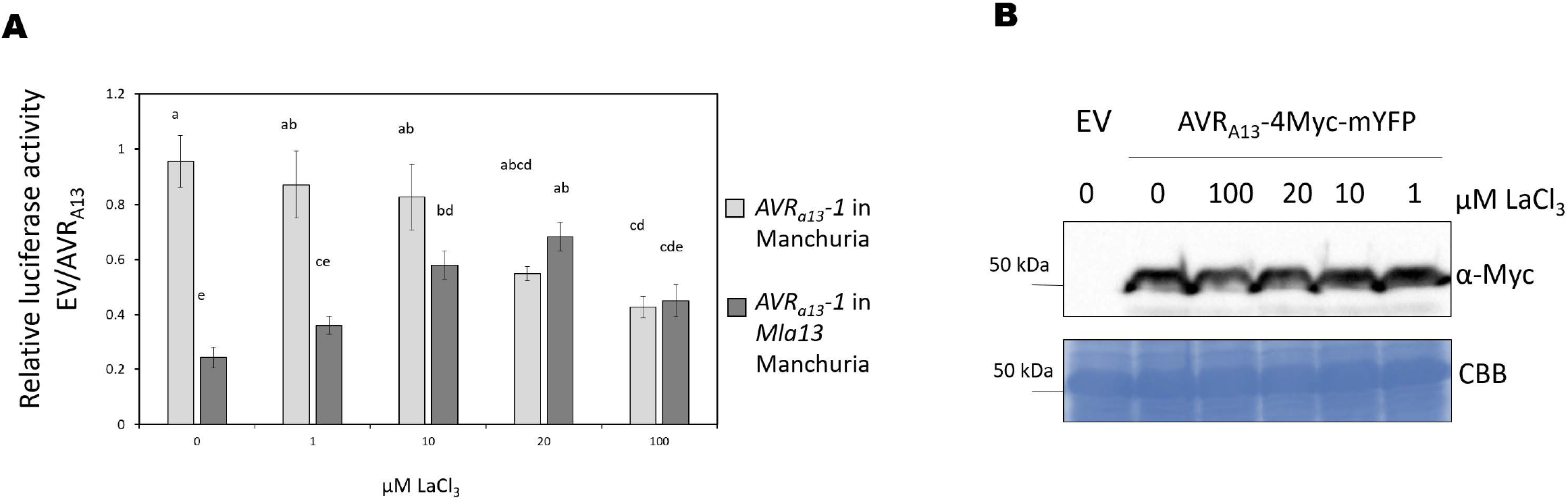
Calcium channel activity is required for *Mla13*-mediated cell death in barley. **(A)** Barley protoplasts of lines CI 16155 (cultivar Manchuria *Mla13)* and CI2330 (Manchuria) were transfected with *pUBQ:luciferase* and piPKb002 containing *AVR_a13_-1* cDNA without signal peptide or a piPKb002 empty vector control and recovered in the presence of LaCl_3_ at concentrations indicated. Luciferase activity was determined 16 hr post transfection/addition of LaCl_3_ as a proxy for cell death and normalized against the respective *EV* sample. Error bars = standard deviation. Differences between samples were assessed using non-parametric Kruskal-Wallis and subsequent Dunn’s post hoc tests. *p*= 6.l79e-lO. Samples marked by identical letters in the plot did not differ significantly (p<0.05) in Dunn’s test. **(B)** Protoplasts derived from cultivar Manchuria CI2330 leaves transfected with *pZmUBQ:AVR_a13_-1-mYFP* were harvested 16h post transfection/LaCl_3_ treatment. Total protein was extracted, separated by gel electrophoresis, and western blots (WB) were probed with anti-GFP antibodies. CBB: Coomassie brilliant blue.

## Discussion

Functional studies of effector recognition by NLRs are not only important for a better understanding of plant disease resistance but also for dissecting the mechanisms pathogens employ to overcome NLR-mediated resistance. To address both aspects, we studied MLA13-mediated recognition of the barley powdery mildew AVRa_A13_ effector family with a particular focus on AVR_A13_-V2, which originated from a *Bgh* isolate that has overcome *Mla13* resistance.

### Mutations in the NB site of MLA13 abrogate association with its matching effector

The residues of the MLA LRR domains, which are under purifying selection, are thought to serve as effector contact residues (Seeholzer *et al*., 2010; Maekawa *et al*., 2019). Residues S^902^ and P^935^ in the 14^th^ and 15^th^ LRRs of MLA13 are exchanged for other amino acids in MLA13 encoded by a cultivar that has lost *Mla13* resistance function (Bettgenhaeuser et al., 2021), and we showed here that the residues are indeed required for effector binding and activation of MLA13. Importantly, however, our data show that not only the contact residues in the NLR LRR domain mediate receptor-effector association, but that an intact, ADP-bound receptor conformation is required for efficient effector-receptor association in yeast. Disruption of this intact conformation by mutations in the NB site of MLA13, which result in the so-called ‘MHD’ (mimicking ATP binding) and ‘P-loop’ (no binding of ADP/ATP) receptor versions (Fig. S4) fully abrogated interaction with the matching AVR_A13_ effector variants in Y2H assay, probably because of spatial hindrance. One possible explanation for this hindrance is that residues of the MLA13 NB domain are engaged in the formation of an effector-accessible conformation of the MLA LRR domain, i.e. a site of effector entry (Förderer *et al*., 2022b) only provided by ADP-bound MLA13 (Fig. S4). At this effector entry site of ADP-bound MLA13, the MLA13 NB domain may transiently contact the AVR_A13_ ligand and this contact may be required for the the steric clash that dislocates the NB domain for exchange of ADP to ATP. In fact, one intermediate state structure of the ADP-bound ZAR1 monomer bound to the activating PBL2 ligand (PDB 6j5v) implies contact between the ZAR1 NB domain and the PBL2 ligand ultimately before the steric clash that allows effector-mediated ZAR1 resistosome formation, although association between the contact-forming residues cannot be detected in the active, ATP-bound ZAR1 resistosome (Wang *et al*., 2019b). An alternative hypothesis of our findings is a transient association between AVR_A13_ and MLA13, implying that conformational changes of MLA13 to the active oligomeric ATP-bound state lead to dislodging of AVR_A13_ effectors from the resistosome complex. However, this model is in contrast with the observation of all active NLR resistosome structures available to date, where each NLR monomer stably binds one activating ligand. The autoactive wheat CNL Sr50^MHD^ mutant was also impaired in AvrSr50 association when compared to wild-type Sr50 (Fig. S3), but our data contrast with the example of enhanced association between the flax TNL L6 MHD version and its matching effector (Bernoux *et al*., 2016). We therefore suggest different requirements for NB domains at the site of effector entry in CNLs and TNLs or for individual NLRs in general. However, we cannot entirely exclude that this difference may be due to the initiation of yeast cell death upon expression of CNL^MHD^, whereas TNL^MHD^ variants cannot induce cell death in yeast. However, the MLA13^MHD^ and Sr50^MHD^ protein levels are as stable as those of wild-type receptors and yeast growth in the presence of leucine is similar between yeasts expressing wild-type receptors and the MHD variants (Fig. 3B and Fig. S3D).

We and others have previously attempted to detect interaction between CNLs and their matching effector *in planta* by using NLR P-loop mutants to prevent NLR-mediated cell death. Blocking TNL ROQ1-mediated cell death signalling in *eds*1 knockout lines in *N. benthamiana* was important for purification of the tetrameric ROQ1-effector resistosome (Martin *et al*., 2020). Our data showing that MLA13 P-loop variants have lost the ability to bind matching effectors might explain why previous attempts to detect effector – receptor interaction using P-loop CNL mutants were unsuccessful.

### Amino acid exchanges in the MLA13 α1-helix deregulate auto-inhibition but not Ca^2+^ - dependent MLA13 cell death function

Negatively charged residues in the α1-helix of NLR CC domains are thought to be required for Ca^2+^ channel activity of CNL resistosomes (Förderer *et al*., 2022b). This was inferred from the observation that replacement of these residues with alanine abrogated ZAR1 Ca^2+^ channel activity and ZAR1-mediated resistance. We observed that the negatively charged residues MLA13D^2^ and MLA13^E17^ in the α1-helix are not required for MLA13-mediated cell death and that these amino acid exchanges instead lead to effector-independent cell death in *N. benthamiana*. We speculate that in the absence of a matching effector, these negatively-charged amino acids in MLA13 are required for burying the α1-helix and that this auto-repression malfunctions in MLA13^D2A_E17A^, i.e. the α1-helix is exposed and available for oligomerization (Fig. S4). However, our data cannot clarify whether the hypothetical auto-active α1-helix conformation of MLA13^D2A_E17A^ allows the exchange of ADP to ATP or whether an ADP-bound NB domain is even capable of forming a functional oligomer (Fig. S4).

The cell death autoactivity of MLA13^D2A_E17A^ contrasts with similar ZAR1 mutants, which abolish cell death, but the data is comparable to results reported for other CNLs, including wheat Sr35 (Adachi *et al*., 2019; Förderer *et al*., 2022a). Despite these differences, we demonstrate that MLA13-dependent and AVR_A13_-triggered cell death activity in barley protoplasts is impaired in the presence of the cation channel inhibitor LaCl_3_, suggesting that cation transport across plant cell membranes by a putative MLA13 channel and/or other cation channels is also an important biochemical activity of the deduced MLA13 resistosome. Although the exact mechanism for cation transport in the putative MLA13 resistosome remains to be determined, our data align with reports on other CNLs that confer calcium channel-dependent cell death (Grant *et al*., 2000; Förderer *et al*., 2022a) and underline that perturbation of Ca^2+^ homeostasis is a fundamental component of both, TNL- and CNL-mediated cell death in plants (Jubic *et al*., 2019; Saur *et al*., 2021; Jacob *et al*., 2021; Förderer *et al*., 2022a).

### A single effector residue can disrupt NLR activation

As LRR domains have the potential to bind a variety of proteinaceous ligands, engineering the LRR domains of NLRs to bind pathogen effectors that are not recognized by the natural immune system appears to be an attractive strategy for controlling plant diseases. Our data demonstrate that ligand binding *per se* is not sufficient for NLR activation and that a single surface-exposed residue, L^119^ in full-length AVR_A13_-V2, can abrogate NLR activation despite enhanced interaction. This dominant-acting interaction may directly allow AVR_A13_-V2 to outcompete all AVR_A13_-1 effectors for association with MLA13 and subsequent receptor activation. Alternatively, AVR_A13_-V2 sequestration of some MLA13 monomers might be sufficient to disrupt putative MLA13 resistosome formation if a threshold of ligand-activated CNLs must be available for pentameric CNL resistosomes to be formed (Förderer *et al*., 2022b). The possibility that AVR_A13_-V2 sequesters AVR_A13_-1 from activation of MLA13 appears less likely because AVR_A13_-V2 can also inhibit MLA13 auto-activity (Fig. 2). The contact residues responsible for the activation of MLA13 by AVR_A13_ are likely unique, despite the overall structural similarity of AVR_A_ effectors and allelic, highly sequence similar MLA receptors (Seeholzer *et al*., 2010; Bauer *et al*., 2021). This appears to be also true for the residues of AVR_A13_-V2 (including L^119^) that mediate MLA13 interaction, as neither the enhanced interaction, nor the dominant-negative effect of AVR_A13_-V2 was detected when MLA13 was replaced by the highly sequence-similar MLA1 or MLA7 NLRs. The overall high sequence and predicted structural identity between AVR_A13_-1 and AVR_A13_-V2, as well as the identification of a single residue, L^119^ of AVR_A13_-V2, as the main driver of enhanced MLA13 interaction, suggest that the binding surfaces to the MLA13 receptor overlap. However, our data implies that AVR_A13_-V2 locks MLA13 into an inactive, effector-bound state by preventing the receptor from transitioning to one of the conformational changes downstream of effector binding (Fig. S5). AVR_A13_-V2 cannot inhibit cell death signalling of MLA13 constitutive-gain of function mutants with amino acid replacements in the CC domain despite interaction with MLA13^D2A_E17A^ (Fig. 4D). We therefore suggest that the inhibitory function of AVR_A13_-V2, mediated by L^199^, affects conformational changes that take place before the release of the MLA13 α1-helix; i.e. AVR_A13_-V2 binding to MLA13 either fails to induce an inter-domain steric clash in the receptor or blocks the transition to the steric clash-mediated open conformation, which allows exchange of ADP to ATP in the NB site of MLA13 (Fig. S5). Alternatively, AVR_A13_-V2 binding to MLA13 induces a steric clash, but AVR_A13_-V2 association inhibits the release of the α1-helix from autorepression. As MLA13 MHD mutants are generally inaccessible to effector binding in Y2H assay (including binding to avirulent AVR_A13_-1), our data cannot clarify whether the loss of inhibitory function of AVR_A13_-V2 on MLA13 cell death takes place before or after ADP exchange to ATP in wild-type MLA13. Collectively, we demonstrate that the stable interaction between AVR_A13_-V2 and inactive MLA13 has the potential to define distinct conformations of intermediate states of CNL receptors. This knowledge is currently largely elusive for both animal and plant NLRs. Understanding such conformations will help ensure that future synthetic NLRs do not become locked into intermediate non-functional states.

### Role of AVR_A13_-V2 in the breakdown of *Mla13* resistance in the European *Bgh* population

Evasion of NLR-mediated pathogen recognition is usually mediated by diversification of the pathogen’s effector repertoire, including allelic variation of effector genes that results in abrogation of effector-NLR receptor associations. This model applies to the virulent variant AVR_A13_-V1. However, AVR_A13_-V2 not only interacts strongly with MLA13, but also inhibits MLA13 cell death signalling in a dominant manner. This raises the possibility that *Bgh* AVR_A13_-V2 facilitates rapid clonal dispersal of virulence in *Bgh* populations that are avirulent on *Mla13*. In the European *Bgh* population the virulence frequency on *Mla13* increased from 0.2% in the 1980s to as high as 60% in 1995 (Gacek, 1987; Jørgensen, J.H.; Hovmøller, 1987; Hovmøller *et al*., 2000), suggesting a major shift in genetic variation of *AVR_a13_* on a continental scale. By contrast, virulence on *Mla13* barley appears to be low at a global scale, with only 7% of *Bgh* isolates in a global strain collection overcoming *Mla13-* mediated resistance (Lu *et al*., 2016; Rsaliyev *et al*., 2017; Saur *et al*., 2019a). In addition, *AVR_a13_/BGH_20990* has a very low frequency of non-synonymous SNPs in tested global and local *Bgh* populations (0.9 non-synonymous SNPs/100 bp coding sequence), indicating an overall low genetic diversity of *AVR_a13_* (Saur *et al*., 2019a). Our data demonstrate a dominant negative activity of *AVR_A13_-V2* on MLA13 therefore suggesting that the breakdown of *Mla13* resistance was caused by direct manipulation of the receptor activation mechanism rather than by evasion of MLA13 recognition.

## Materials and methods

### Plant and fungal materials and growth conditions

Near isogenic lines (NILs) of the barley cultivar Manchuria were grown at 19 °C, 70% relative humidity, and under a 16 h photoperiod. *N. benthamiana* plants were grown under standard greenhouse conditions under a 16 h photoperiod. Maintenance of *Bgh* isolates was carried out as described previously (Lu *et al*., 2016).

### Generation of expression constructs

For transient gene expression assays in *N. benthamiana* and barley protoplasts and for yeast 2-hybrid interaction studies, coding sequences of receptor and effector genes with or without stop codons were either synthesized as pDONR221 entry clones from GeneArt (Thermo Scientific), or were published previously (Saur *et al*., 2019a). Respective genes were transferred from entry or donor vectors into the expression vectors pIPKb002 (Himmelbach *et al*., 2007), pGWB414, pGWB517 (Nakagawa *et al*., 2007), pXCSG-GW-HA, pXCSG-GW-Myc, pXCSG-GW-mYFP (Garcia *et al*., 2010), pAMpAT-GW-BirA-4Myc, pLexA-GW, or pB42AD-GW (Shen *et al*., 2007) as indicated using LR Clonase II (Thermo Scientific).

### Transient gene expression by Agrobacterium-mediated transformation of Nicotiana benthamiana leaves

*Agrobacterium tumefaciens* GV3101:pMP90K were freshly transformed with respective constructs of interest and grown from single colonies in liquid Luria broth medium containing appropriate antibiotics for ~ 24 hours at 28 °C to an OD_600_ not higher than 1.5. Bacterial cells were harvested by centrifugation at 2500 *× g* for 15 min followed by resuspension in infiltration medium (10 mM MES, pH 5.6, 10 mM MgCl_2_, and 200 μM acetosyringone) to a final OD_600_ = 1. Cultures were incubated for two to four h at 28 °C with 180 rpm shaking before infiltration into leaves from three to five-week-old *N. benthamiana* plants. For co-expression of multiple constructs, Agrobacteria carrying the genes of interest were mixed equally unless indicated otherwise. Cell death was assessed one to five days post infiltration as indicated and tissue for immunodetection analysis was harvested one to two days post infiltration as indicated.

### Protein extraction from Nicotiana benthamiana leaf tissue for protein detection by immunoblotting

Frozen leaf material was ground to a fine powder using pre-cooled adapters in a bead beater (Retsch) and thawed in cold plant protein extraction buffer (150 mM Tris-HCl, pH 7.5, 150 mM NaCl, 10 mM EDTA, 10% (v/v) glycerol, 5 mM DTT, 2% (v/v) plant protease inhibitor cocktail (Sigma), 1 mM PMSF, and 0.5 % (v/v) IGEPAL) at a ratio of 50 mg fresh tissue/150 μl of extraction buffer. Extracts were centrifuged twice at 15,000 *× g* for 10 min at 4 °C. For SDS-PAGE, extracts were diluted 4:1 with 4x SDS loading buffer and heated to 85 °C for 10 to 15 min before again removing insoluble material by centrifugation at 15,000 *× g* for 5 min. For pull-down of mYFP-tagged proteins, GFP-Trap-MA (Chromotek) beads were incubated in equilibration buffer for 1 h at 4 °C and subsequently mixed with one ml of protein extracts for 2 to 3 h at 4 °C with slow but constant rotation. Then, conjugated GFP-Trap beads were washed five times in 1 ml of cold wash buffer at 4 °C before interacting proteins were stripped from the beads by boiling in 25 μl of 4x SDS loading buffer for 5 min. Samples were separated on 8% to 13% SDS-PAGE gels, blotted onto PVDF membrane, and probed with anti-GFP (abcam ab6556), anti-Myc (abcam ab9106) or anti-HA (Roche 3F10) followed by anti-rabbit IgG-HRP (Santa Cruz Biotechnology sc-2313) or anti-rat IgG-HRP (abcam ab97057) secondary antibodies. Epitope-tagged proteins were detected by the HRP activity on SuperSignal West Femto Maximum Sensitivity Substrate (Thermo Fisher 34095) using a Gel Doc™ XR+ Gel Documentation System (Bio-Rad).

### Proximity-dependent protein labelling of proteins transiently expressed in Nicotiana benthamiana leaves

Pull-down of biotinylated proteins was performed by following published protocols (Conlan *et al*., 2018) with the alteration that free biotin was not removed before adding streptavidin to protein extracts. Instead, we applied a 10 μM biotin solution to the plant tissue (instead of a 75 μM solution (Conlan et al., 2018). We followed a sequence of infiltrations to minimise MLA-mediated cell death of *N. benhamiana* leaf tissue: *Agrobacterium tumefaciens* GV3101::pMP90K carrying *35S:Mla-4Myc* constructs were grown from glycerol stocks and infiltrated (day 1). At 24 h post infiltration of the *Mla* constructs, Agrobacteria freshly transformed with *35S:AVR_a13_-BirA-4Myc* constructs or *EV* were infiltrated as indicated (day 2). Ten μM of free biotin in infiltration buffer lacking acetosyringone was infiltrated at 24 h after the second infiltration and 48 h after the first infiltration (day 3). Tissue for streptavidin-based precipitation of biotinylated proteins was harvested 24 h post infiltration of free biotin. Frozen leaf material was ground to a fine powder using pre-cooled adapters in a bead beater (Retsch) and thawed in cold plant denaturing extraction buffer (150 mM Tris-HCl, pH 7.5, 150 mM NaCl, 10 mM EDTA, 5% (v/v) glycerol, 5 mM DTT, 1% (v/v) plant protease inhibitor cocktail (Sigma), 1 mM NaF, 1 mM sodium orthovanadate, 1 mM PMSF, 1% TritonX-100 and 0.5 % (w/v) SDS) at a ratio of 300 mg fresh tissue/2 ml of denaturing extraction buffer. Extracts were incubated rotating at 4°C for 30 minutes before the removal of insoluble material by centrifugation at 21,000 × g for 30 min at 4 °C. Streptavidin coated Dynabeads (100 μL/sample, MyOne streptavidin C1, Thermo Fisher) were incubated in wash buffer (150 mM Tris-HCl, pH 7.5, 150 mM NaCl, 10 mM EDTA, 5% (v/v) glycerol, 1% (v/v) plant protease inhibitor cocktail (Sigma)) containing 1% BSA for 1 h at 4 °C and subsequently mixed with 2two ml of protein extracts for 3 h at 4 °C with slow but constant rotation. Then, conjugated Streptavidin beads were washed four times in 1 ml of cold wash buffer before interacting proteins were stripped from the beads by heating to 85°C for 10-15 min in 50 μl of 4x SDS loading buffer. From these 50 μl, 30μl were loaded on 9% SDS-PAGE gels. Proteins were blotted onto PVDF membrane and probed with anti-Myc (abcam ab9106) followed by anti-rabbit IgG-HRP (Santa Cruz Biotechnology sc-2313) secondary antibodies. Myc-tagged proteins were detected by the HRP activity on SuperSignal West Femto Maximum Sensitivity Substrate (Thermo Fisher 34095) using a Gel Doc™ XR+ Gel Documentation System (Bio-Rad).

### Transient gene expression and cell death assay in barley protoplasts

Assessment of protoplast cell death using a luciferase activity as a proxy for cell viability was performed as described (Saur *et al*., 2019b). Briefly, *AVR_a13_ -V2* cDNA lacking the respective signal peptide was expressed from the Zea mays ubiquitin promotor in protoplasts isolated from barley cultivar Manchuria CI 2330 and cultivar Manchuria *Mla13* NIL CI 16155. For this, the epidermis of the primary leaves from seven to eight-day-old plants was removed before leaves were immersed in the enzyme solution. A total volume of 30 μl water containing 5 μg of the *luciferase* reporter and 6 μg of the *AVR_a13_-V2* effector construct or an *EV* was transfected into 300 μL barley protoplasts at a concentration of 5 *×* 10^5^ protoplasts/ml solution. Protoplasts were recovered in regeneration buffer supplemented with LaCl_3_ as indicated. About 16 h after transfection, protoplasts were collected by centrifugation at 1000 *× g*, the supernatant was discarded, and 200 μl 2x cell culture lysis buffer were added (Promega, E1531). Luciferase activity was determined by mixing 50 μl of protoplast lysate with 50 μl luciferase substrate (Promega, E1501) in a white 96-well plate and light emission was measured at 1 second/well using a microplate luminometer (Centro, LB960).

### Protein extraction from barley protoplasts and fusion protein detection by immunoblotting

To determine the effect of LaCl_3_ treatment on AVR_A13_ protein, for each LaCl_3_ treatment, 300 μg of the *AVR_a13_-V2-mYFP* effector construct or an *EV* was transfected into 3 ml pf barley protoplasts cultivar Manchuria CI 2330 at a concentration of 5 *×* 10^5^ protoplasts/ml solution. Protoplasts were recovered in regeneration buffer supplemented with the LaCl_3_ to the final concentrations indicated. About 16 h post transfection, protoplasts were collected by centrifugation at 1000 *× g*, the supernatant was discarded and protoplast pellets were frozen in liquid nitrogen. Total protein was extracted by the addition of 100 μl cold plant protein extraction buffer (200 mM Tris-HCl, pH 7.5, 150 mM NaCl, 10 mM EDTA, 10% (v/v) glycerol, 12 mM DTT, 2% (v/v) plant protease inhibitor cocktail (Sigma), and 1 % (v/v) IGEPAL) to each protoplast pellet. Extracts were centrifuged at 15,000 *× g* for 5 min at 4 °C. For SDS-PAGE, extracts were diluted 4:1 with 4x SDS loading buffer and heated to 85 °C for 10 to 15 min before removing insoluble material by centrifugation at top speed for 5 minutes. Samples were separated on 10% SDS-PAGE gels, blotted onto PVDF membrane, and probed with anti-GFP (Santa Cruz Biotechnology sc-8334 or abcam ab6556) followed by anti-rabbit IgG-HRP (Santa Cruz Biotechnology sc-2313) secondary antibodies. mYFP tagged proteins were detected by the HRP activity on SuperSignal West Femto Maximum Sensitivity Substrate (Thermo Fisher 34095) using a Gel Doc™ XR+ Gel Documentation System (Bio-Rad).

### Yeast 2-hybrid assay and yeast protein extraction

*NLR* receptor gene variants were cloned into the pLexA-GW vector (Shen *et al*., 2007) for expression with an N-terminal LexA activation domain under the control of a constitutive ADH1 promoter (BD-NLR). Effector variants were cloned into pB42AD-GW (Shen *et al*., 2007) for expression with an N-terminal B42 activation domain followed by the HA-tag under the control of an inducible GAL_1_ promoter (AD-AVR). Using the lithium acetate method (Gietz and Woods, 2002), bait and prey constructs were co-transformed into the yeast strain EGY4.8 p8op and successful transformants were selected by colony growth on SD-UHW/Glu (2% (w/v) Glucose, 0.139% (w/v) yeast synthetic drop-out medium pH 6 without uracil, histidine, tryptophan, 0.67% (w/v) BD Difco yeast nitrogen base, 2% (w/v) Bacto Agar). Yeast transformants were grown to OD_600_ = 1 in liquid SD-UHW/Glu before harvesting cells for drop out of the dilution series on SD-UHW/Gal/Raf media (SD-UHW without glucose but with 2% (w/v) Galactose 1 % (w/v) Raffinose, with (-UHW) or without Leucine (-UHWL)) and incubated for one to two weeks at 30 °C.

For protein detection, yeast strains were grown to OD_600_ = 1 in SD-UHW/Gal/Raf liquid medium at 30 °C and 200 rpm shaking, and proteins were extracted using 200 mM NaOH (NaOH method) (Zhang *et al*., 2011)). Total protein samples were separated on 9% or 12% SDS-PAGE gels, blotted onto PVDF membrane, and probed with anti-HA (Merck, clone 3F10) or anti-LexA (Santa Cruz Biotechnology, sc7544) primary antibodies followed by anti-rat (Santa Cruz Biotechnology, sc2065) or anti-mouse IgG-HRP (Santa Cruz Biotechnology, sc2005) secondary antibodies as appropriate. HA and LexA fusion proteins were detected by HRP activity on SuperSignal West Femto Maximum Sensitivity Substrate (Thermo Fisher 34095) using a Gel Doc™ XR+ Gel Documentation System (Bio-Rad).

## Supplement

**Figure S1:** Proximity-dependent protein labelling confirms requirement of AVR_A13_ C-terminus for MLA13 interaction.

**Figure S2:** Specificity control to Fig. 1D.

**Figure S3:** Gain-of-function and loss-of-function NLR mutants and their ability to bind matching avirulence effectors.

**Figure S4:** Schematic model of MLA13 wild-type and mutant conformations.

**Figure S5:** Schematic models of MLA13 activation by *Bgh* AVR_A13_ −1 and inhibition by AVR_A13_-V2,respectively.

**Supplemental Raw Data:** raw data of all figures.

## Acknowledgements

We would like to thank Sabine Haigis and Petra Köchner for technical support and maintenance of *Bgh* isolates and Ksenia Krasileva for critical comments on the manuscript. Although the resulting crosses could not be assessed for *Bgh* infection due to loss of MLA13 resistance in control lines, we highly acknowledge the group of Matthew Moscou that crossed *Mla13* barley with *AVR_a13_-V2* transgenic lines for assessing AVR_A13_-V2-mediated inhibition of Mla13 resistance. IMLS, TM and PSL acknowledge support from the Cluster of Excellence on Plant Sciences (CEPLAS) funded by the Deutsche Forschungsgemeinschaft (DFG, German Research Foundation) under Germany’s Excellence Strategy – EXC 2048/1 – Project ID: 390686111 and funding from the DFG Collaborative Research Centre Project-ID 414786233 (SFB 1403). This work was also funded by the DFG Emmy Noether Programme (SA 4093/1-1 to IMLS), the Daimler and Benz Foundation (IMLS) and the Max Planck Society (PSL).

## Author contributions

EEC, MBS, TM, PSL & IMLS designed the research, EEC, MBS and IMLS performed the experiments and data analysis. EEC, IMLS and PSL wrote the paper with contributions from all authors.

## Data availability

All relevant data are available within the paper and its supplementary data published online.

## Figures

**Figure S1:**
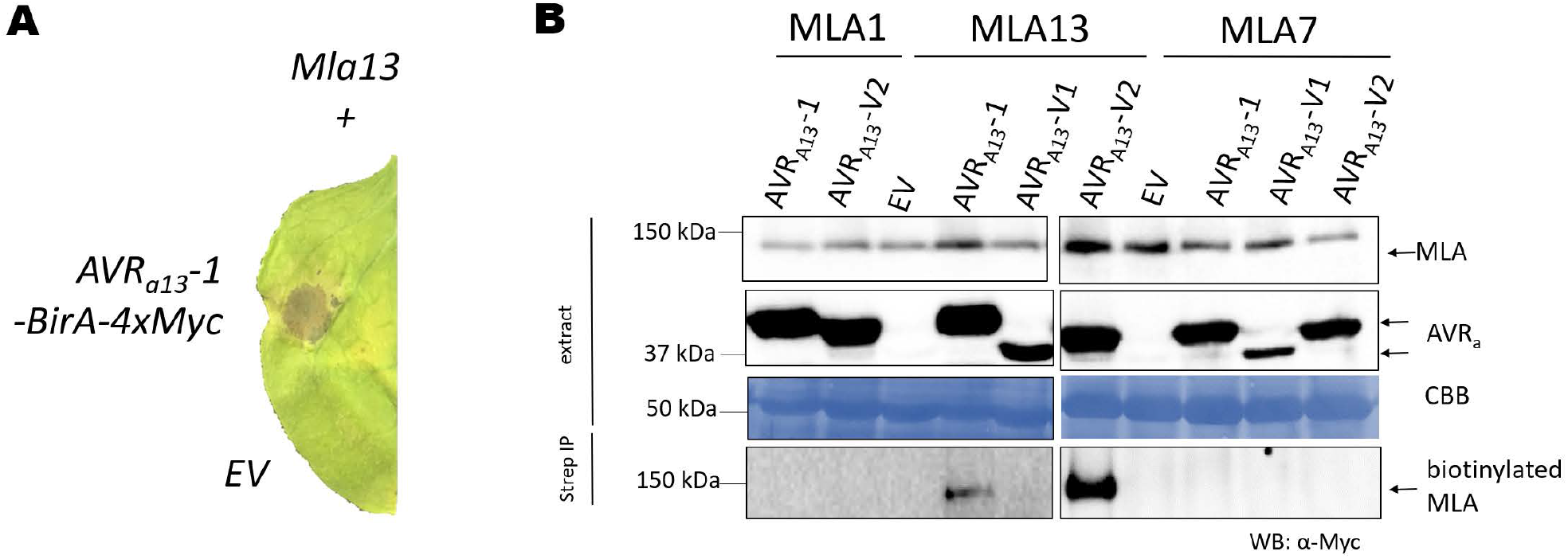
**(A)** *Nicotiana benthamiana* leaves were transformed transiently with cDNAs of the *Mla13* together with *empty vector (EV)* or *AVR_a13_-1* lacking SPs and fused c-terminally to *BirA-4Myc* tag sequence and expressed from the 35S promotor. Cell death was determined three days post transformation and picture shows representative of at least three independent leaves. **(B)** *N. benthamiana* leaves were transformed transiently with cDNAs of *Mla1* or *Ml_a7_* or *MLA13* fused C-terminally to a *4xMyc* sequence and at 24 h before re-transformation with cDNAs encoding *AVR_a13_-1-BirA-4xMyc, AVR_a13_-V1-BirA-4xMyc, AVR_a13_-V2-BirA-4xMyc* or *empty vector (EV)* as indicated. All leaves were treated with 10 μM biotin by infiltration at 24h after the second transformation. Leaf tissue was harvested 24h post biotin treatment. Total protein was extracted under denaturing conditions and recovered by Strep IP, separated by gel electrophoresis and probed by anti-Myc western blotting (WB). CBB: Coomassie brilliant blue.

**Fig. S2:**
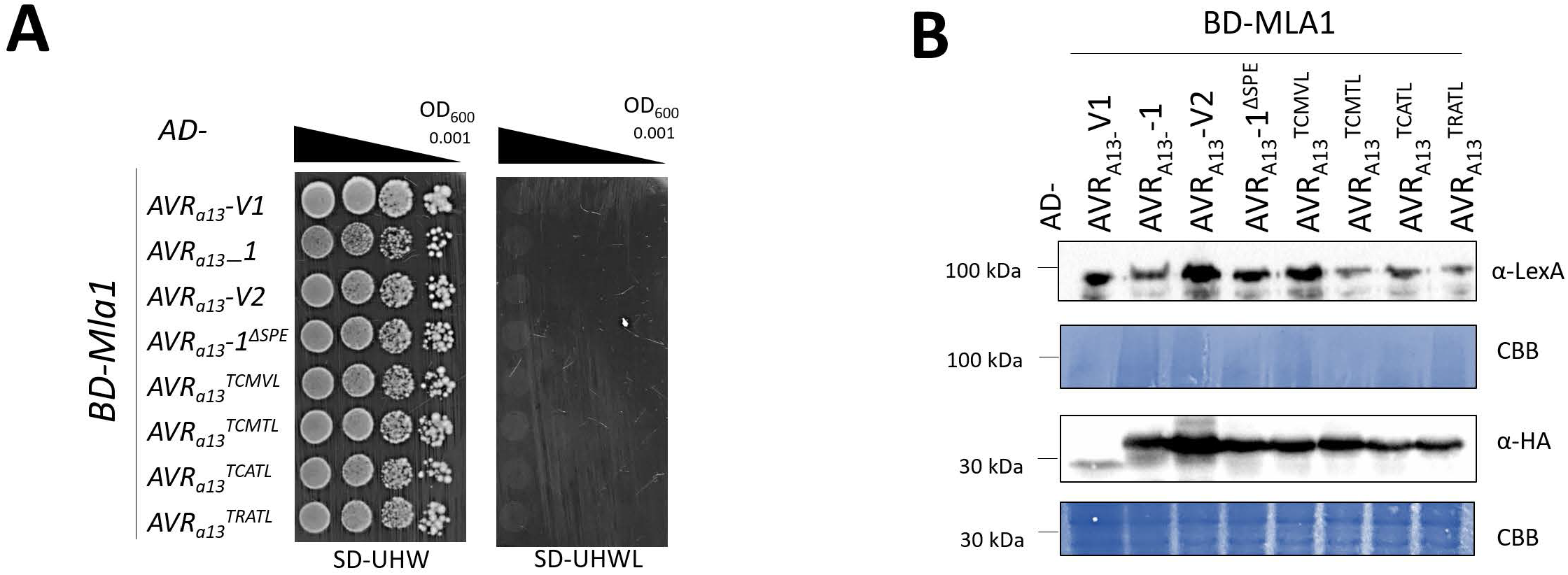
Specificity control to Figure 1D. **(A,B)** Yeast cells were co-transformed with *Mla1* fused N-terminally to the *LexA* binding domain sequence (BD) and *AVR_a13_* variants lacking SPs fused N-terminally to the *B42* activation domain (AD) and 1xHA tag sequence as indicated. Growth of transformants was determined on selective growth media containing raffinose and galactose as carbon sources but lacking uracil, histidine and tryptophan (-UHW), and interaction of proteins was determined by leucine reporter activity reflected by growth of yeast on selective media containing raffinose and galactose as carbon sources but lacking uracil, histidine, tryptophan and leucine (-UHWL). Figures shown are representatives of at least three experiments and pictures were taken 6 to 8 days after drop out. (B) Protein levels of BD-MLA1 and AD-AVR_A_ variants corresponding to yeast of D. Yeast transformants were grown in raffinose and galactose containing selective media lacking uracil, tryptophan, and histidine to OD_600_ = 1. Then, cells were harvested, total protein extracted, separated by gel electrophoresis, and western blots (WB) were probed with anti-LexA or anti-HA antibodies as indicated. CBB: Coomassie brilliant blue.

**Figure S3:**
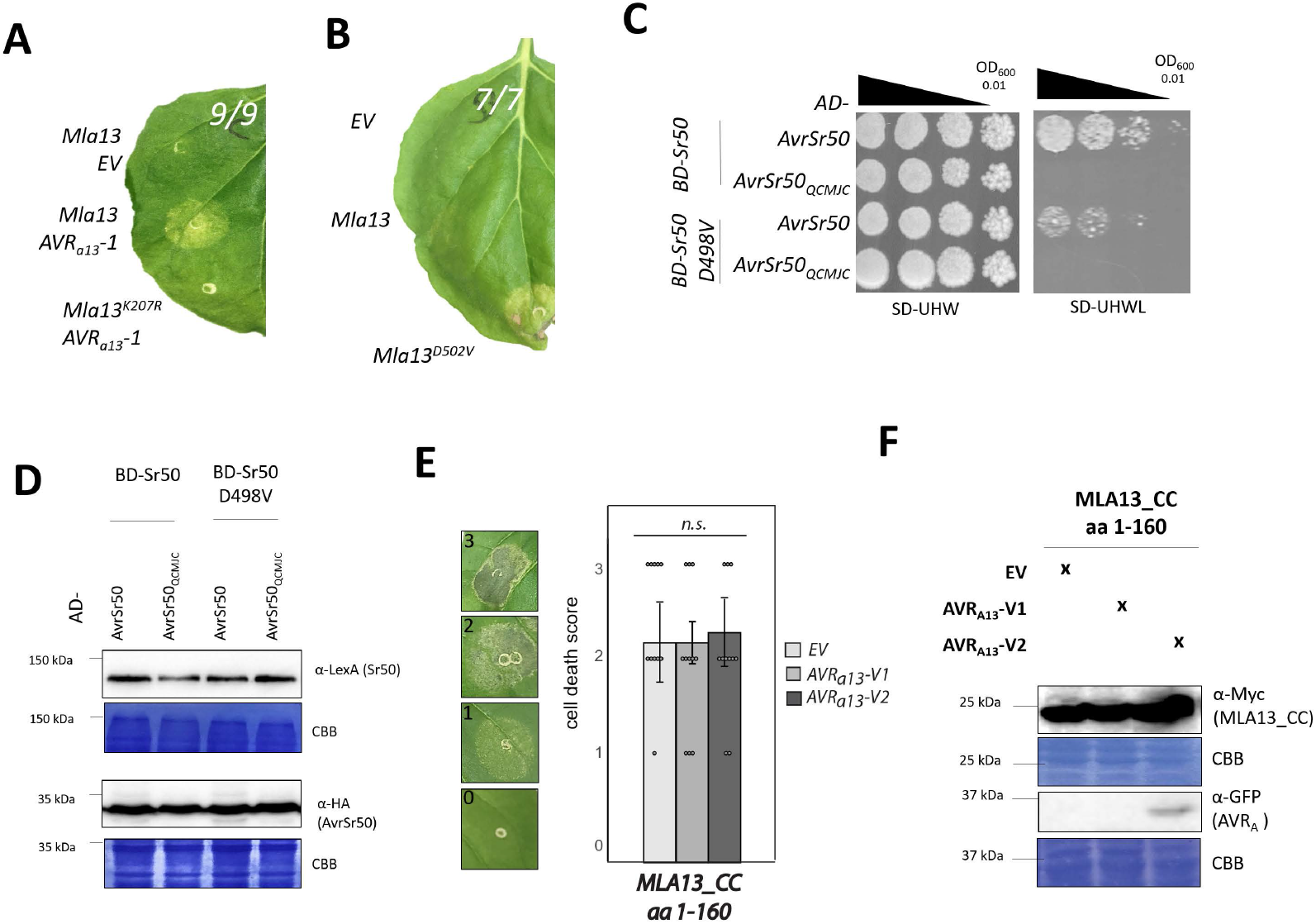
**(A, B)** *Nicotiana benthamiana* leaves were co-transformed transiently with *empty vector (EV)* or constructs encoding either MLA13, MLA13^K207D^ or MLA13^D502V^ (pGWB) as indicated with (A) or without (B) cDNA encoding of *AVR_a13_-1* or *EV*. All cDNAs were under the control of the 35S promoter sequence. Cell death was determined three days post transformation. **(C, D)** Yeast cells were cotransformed with Sr50 or Sr50^D498V^ (MHD) fused N-terminally to the *LexA* binding domain sequence (BD) and AvrSr50 variants lacking SPs fused N-terminally to the *B42* activation domain (AD) and 1xHA tag sequence as indicated. Growth of transformants was determined on selective growth media containing raffinose and galactose as carbon sources but lacking uracil, histidine and tryptophan (-UHW), and interaction of proteins was determined by leucine reporter activity reflected by growth of yeast on selective media containing raffinose and galactose as carbon sources but lacking uracil, histidine, tryptophan and leucine (-UHWL). Figures shown are representatives of at least three experiments and pictures were taken 12 to 14 days after drop out. (B) Protein levels of BD-Sr50 and AD-AvrSr50 variants corresponding to yeast of C. Yeast transformants were grown in raffinose and galactose containing selective media lacking uracil, tryptophan, and histidine to OD_600_ = 1. Then, cells were harvested, total protein extracted, separated by gel electrophoresis, and western blots (WB) were probed with anti-LexA or anti-HA antibodies as indicated. **(E)** *Nicotiana benthamiana* leaves were co-transformed transiently with cDNAs of *AVR_a13_-V1* or *AVR_a13_-V2* or *empty vector (EV)* together with constructs encoding the MLA13 coiled-coil (CC) domain (amino acids (aa) 1-160). **(E)** Cell death was determined two days post transformation and scored from 0 to 3 based on the cell death scale indicated. All values obtained in at least three independent experiments are indicated by dots, error bars = standard error. Differences between samples were assessed by the non-parametric Kruskal-Wallis test. *p*= 0.623871; n.s. = not significant. **(F)** Protein levels corresponding to samples of C. Leaf tissue was harvested 36 hours post infiltration. Total protein was extracted, separated by gel electrophoresis and probed by anti-Myc (MLA13_CC) or anti-GFP (AVR_A13_-V2) western blotting as indicated. CBB: Coomassie brilliant blue.

**Fig. S4:**
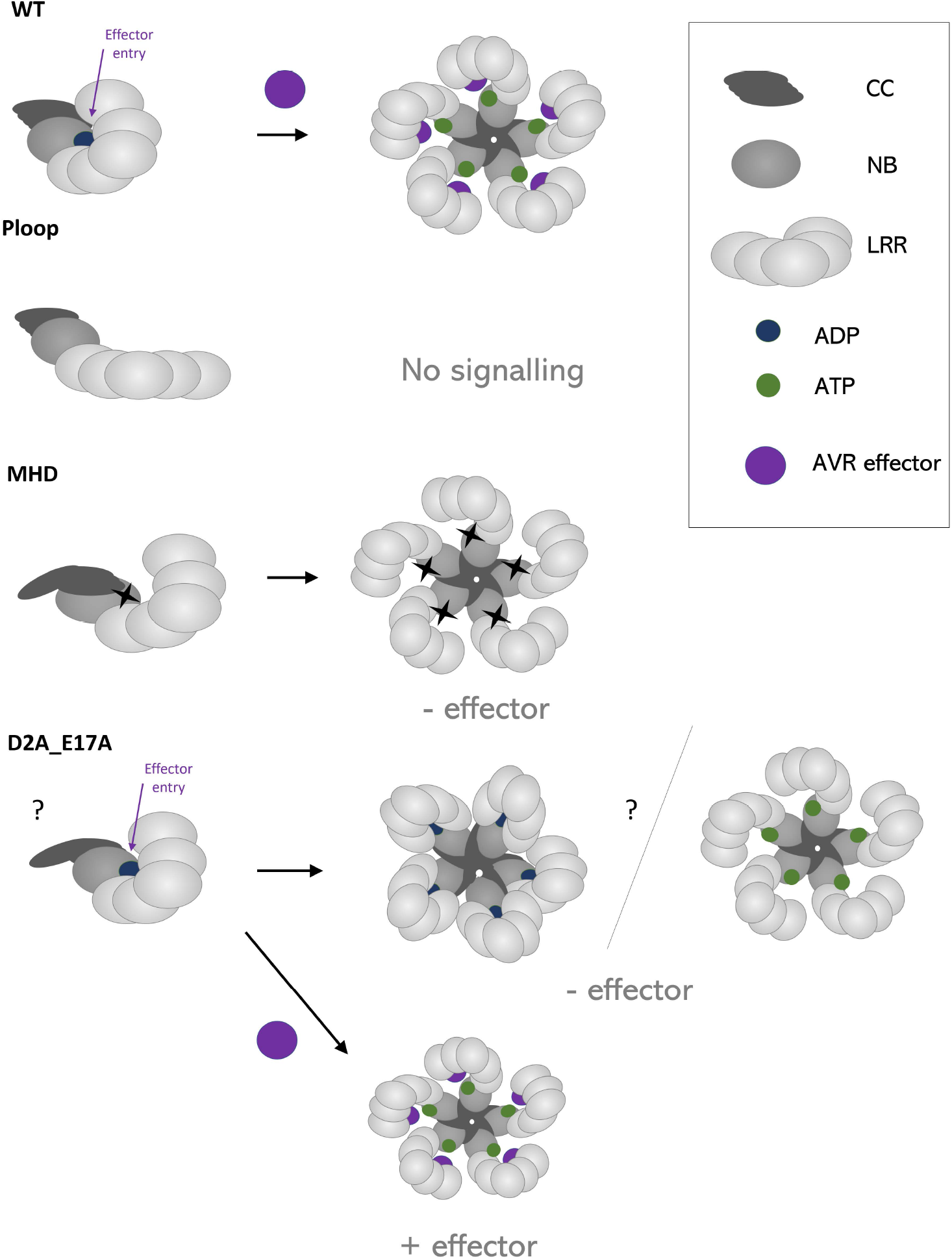
Schematic models of monomeric and oligomeric MLA13 wild-type, MLA13^P-loop^, MLA13^MHD^ and MLA13^D2A_E17A^ conformations with indication of putative effector (purple) entry sites and binding of ADP (green) or ATP (blue).

**Fig. S5:**
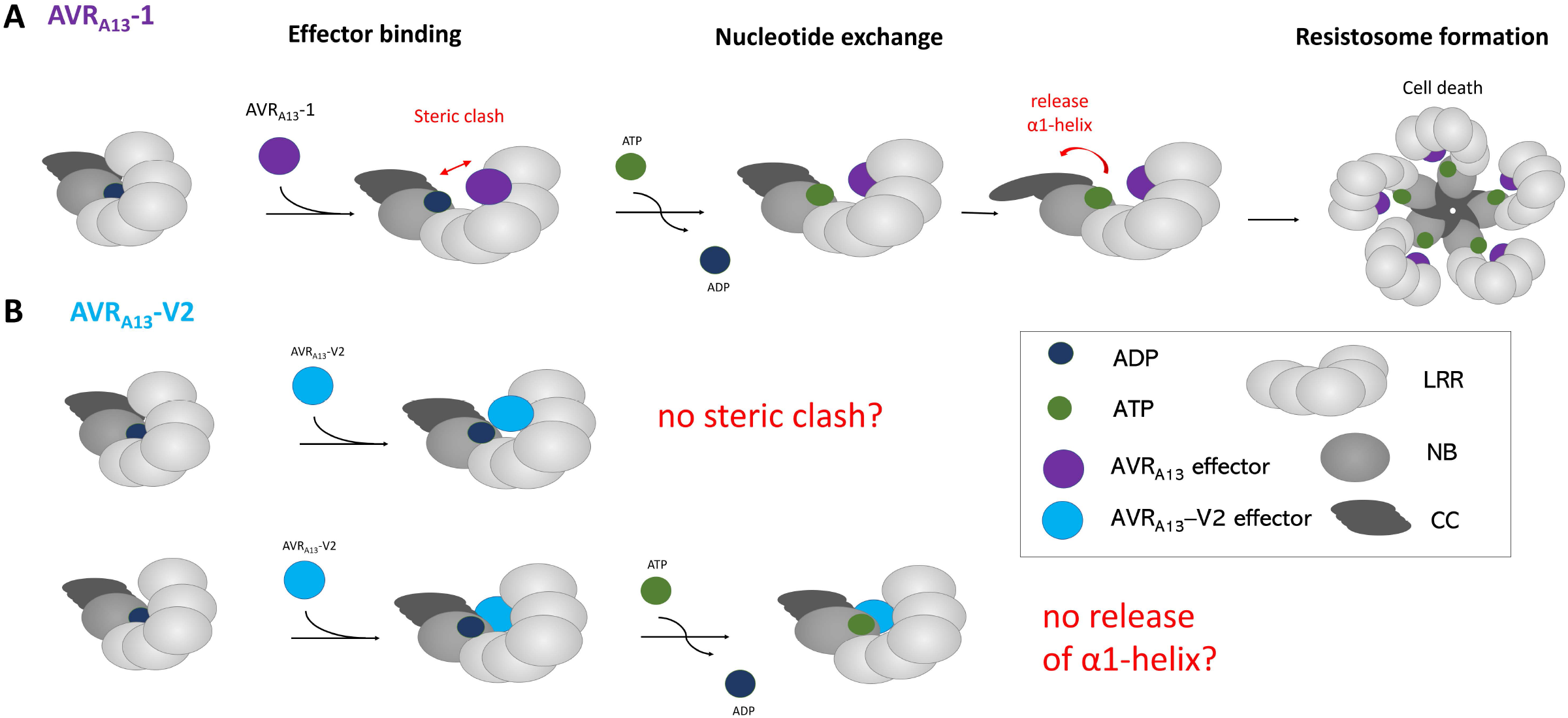
Schematic models of MLA13 during the multistep process of putative resistosome formation initiated by the interaction with *Bgh* AVR_A13_-1 (A) and putative models for the inhibition of the activation process by AVR_A13_-V2 (B). (A) AVR_A13_-1 binding to the effector entry point involving the MLA13 Leucine-rich-repeats (LRR) domain leads to a steric clash and subsequent replacement of adenosine diphosphate (ADP) by adenosine triphosphate ATP) in the nucleotide-binding (NB) pocket of the MLA13. ATP-binding causes additional structural rearrangement of the N-terminal Coiled-coil (CC) domain releasing the α1-helix. In the resulting putative pentameric wheel-like MLA13 resistosome, the α1-helices are thought to form a funnel like structure. (B) AVR_A13_-V2 binding is likely either incapable to inducing a steric clash or prevents subsequent release of the α1-helix.

